# VISUAL EXPERIENCE SHAPES ARM POSITION SENSE IN INTERNAL AND EXTERNAL REFERENCE FRAMES AND ASSOCIATED CORTICAL LOAD

**DOI:** 10.64898/2026.06.08.730866

**Authors:** Kyunggeune Oh, Nikhilesh Natraj, Lewis A. Wheaton, Boris I. Prilutsky

**Affiliations:** School of Biological Sciences, Georgia Institute of Technology, 555 14th Street NW, Atlanta, GA 30332-0356, USA; Department of Mechanical and Measurement & Control Engineering, Idaho State University, 921 S. 8th Ave, Pocatello, ID 83209-8060, USA; Department of Neurology, Weill Institute for Neurosciences, University of California San Francisco, San Francisco, CA 94110, USA

**Author notes:** equal contributions (Kyunggeune Oh, Nikhilesh Natraj). Corresponding authors: Boris I. Prilutsky and Lewis A. Wheaton School of Biological Sciences Georgia Institute of Technology 555 14th Street NW Atlanta, GA 30332-0356, USA.

**Keywords:** Hand position sense, visual experience, precision, accuracy, contingent negative variation

## Abstract

The ability to accurately perceive arm position is essential for motor control and depends on the integration of proprioceptive and visual information. However, how lifelong visual impairment (VI) affects position sense and its neural correlates remains unclear. We quantified arm position sense and associated cognitive-motor load in right-handed visually impaired (n = 7) and normally sighted (NS; n = 7) individuals using three bilateral arm position matching tasks: joint angle matching (JAM; internal coordinates), hand direction–distance matching (DDM; external coordinates), and mirror direction–distance matching (MDDM; external coordinates kinematically identical to JAM). Cognitive load was assessed using the contingent negative variation (CNV) from EEG recordings. VI participants exhibited reduced accuracy and precision of arm position sense in most conditions, and greater CNV magnitude, particularly in the left parietal cortex. Across both groups, performance was worse and CNV magnitude was greater in the DDM task compared with JAM, whereas JAM and MDDM yielded similar behavioral and neural outcomes. These findings indicate that (i) visual experience enhances arm position sense, and (ii) representing limb position in external coordinates imposes greater cognitive demands than encoding joint-based posture. The similarity between JAM and MDDM suggests that participants preferentially rely on internal representations when task kinematics permit.

## INTRODUCTION

Arm position sense—the ability to perceive the location and orientation of individual limb segments—is fundamental for dexterous interaction and efficient navigation within the environment. This sense arises from the complex integration of proprioceptive, cutaneous, and visual inputs developed through childhood and refined in adulthood. Proprioceptive signals underlying limb position sense originate from muscle spindles (Proske and Gandevia, 2018; Banks et al., 2021) and cutaneous afferents from the skin overlying joints (Cohen et al., 1994; Edin, 2001; Aimonetti et al., 2007). These signals are transmitted via the dorsal column and dorsolateral tracts through the medulla and thalamus to the primary somatosensory cortex (Delhaye et al., 2018).

This afferent information defines a joint-based, internal representation of arm position, such as the angles of the elbow and shoulder. However, these signals also serve as the foundation for generating a body-based, external representation of the limb. This process requires estimates of segment dimensions obtained from exteroceptive inputs and a nonlinear coordinate transformation from internal to external reference frames (Soechting and Terzuolo, 1986; Rossetti et al., 1995; Graziano et al., 2000; Oh and Prilutsky, 2022). Current evidence suggests this coordinate transformation is mediated by the posterior parietal cortex (PPC), with the left PPC playing a specialized role in spatial mapping and motor planning, particularly in right-handed individuals (Buneo et al., 2002; Makin et al., 2007; Delhaye et al., 2018).

Beyond providing real-time spatial data, visual experience is vital for the initial formation of somatosensory representations (Medina et al., 2010; Longo et al., 2015). Vision acts as a “teaching signal” that allows the brain to calibrate and refine the body schema in an internal frame of reference (Thinus-Blanc and Gaunet, 1997; Ramachandran and Hirstein, 1998; Crollen and Collignon, 2012). For instance, magnifying the visual image of one’s arm can enhance tactile acuity (Kennett et al., 2001), whereas visual distortions through prisms induce immediate errors in position sense (Mon-Williams et al., 1997). Consequently, individuals with visual impairment (VI) resolve sensory conflicts differently than the normally sighted (NS) individuals (Nava et al., 2014). A lack of visual experience, especially early in life, may impair how proprioceptive information is mapped onto external space (Fiehler et al., 2009; Cappagli et al., 2017).

While compromised vision may negatively affect external spatial perception, some studies suggest that the visually impaired may develop heightened sensitivity to remaining cues through cross-modal plasticity, potentially resulting in more precise somatosensory perception (Goldreich and Kanics, 2003; Pascual-Leone et al., 2005).

The neural cost of performing the internal-to-external coordinate transformations remains poorly understood. In this study, we utilized the Contingent Negative Variation (CNV) potential, i.e. an electrophysiological marker of cognitive-motor load (Walter et al., 1964; Cui et al., 2000), to quantify the neural resources required for arm position matching in internal joint-based and external coordinate frame. We hypothesized that the CNV would reflect the increased preparatory effort needed to transform a proprioceptive percept from one reference frame to another.

We tested two hypotheses: (1) Visual experience enhances arm position sense in external coordinates; and (2) Perceiving arm position in external coordinates is less accurate and more cognitively demanding than arm position perception in internal joint-based coordinates, particularly in visually impaired individuals.

To test these hypotheses, we compared the accuracy and precision of arm position sense between VI and NS individuals across internal joint-based and external reference frames while simultaneously recording 64-channel EEG to assess the cortical demands of these spatial representations.

## METHODS

### Participants

All participants in this study were right-handed (determined based on the Edinburgh Handedness Inventory (Oldfield 1971), over the age of 18 years and had no history of known neurological or musculoskeletal disorders. We recruited seven individuals with normal vision (6 males and 1 female; age = 30.1±13.7 years; mean±SD) and seven age-matched VI individuals (4 males and 3 females; age = 39.3±15.1 years; Table 1). The NS participants participated in another study (Oh and Prilutsky, 2022) that partially presented results of this study. The VI participants were legally blind and had best-corrected visual acuity at or below 20/200 in both eyes according to the Snellen acuity measure (Colenbrander, 1994). Four VI participants were congenitally blind, one participant had been blind since the age of three, and the remaining two participants became blind at the age of 15 and 25 (Table 1). On average, prior to the study, the VI participants had been blind for 33.1±10.7 years.

**Table 1.** Characteristics of participants.

| Subject | Vision | Time with-<br>out vision<br>prior to<br>study (years) | Cause of blindness | Age<br>(years) | Sex | Height<br>(cm) | Forearm<br>+ hand<br>length<br>(cm) | Upper<br>arm<br>length<br>(cm) |
| --- | --- | --- | --- | --- | --- | --- | --- | --- |
| Visually impaired group |  |  |  |  |  |  |  |  |
| 1 | Impaired | 38 | Congenital malfor-<br>mation | 38 | M | 160 | 45.5 | 25.4 |
| 2 | Impaired | 35 | Congenital malfor-<br>mation | 35 | M | 175 | 43.9 | 33.3 |
| 3 | Impaired | 43 | Degenerative retinitis<br>pigmentosa | 58 | M | 180 | 49.3 | 33.5 |
| 4 | Impaired | 31 | Diabetes | 56 | M | 168 | 40.8 | 31.3 |
| 5 | Impaired | 46 | Congenital malfor-<br>mation | 46 | F | 152 | 37.3 | 23.8 |
| 6 | Impaired | 23 | Optic nerve hypoplasia | 23 | F | 152 | 39.1 | 26.4 |
| 7 | Impaired | 16 | Retinoblastoma | 19 | F | 150 | 34.2 | 22.9 |
| Mean |  | 33.1 |  | 39.3 |  | 162.4 | 41.44 | 28.09 |
| ± SD | - | ±10.7 | - | ±15.1 | - | ±12.1 | ±5.16 | ±4.51 |
| Normally sighted group |  |  |  |  |  |  |  |  |
| 1 | Normal | - | - | 19 | M | 165 | 39.83 | 29.25 |
| 2 | Normal | - | - | 47 | M | 178 | 43.69 | 31.88 |
| 3 | Normal | - | - | 29 | F | 156 | 38.94 | 32.12 |
| 4 | Normal | - | - | 20 | M | 175 | 43.74 | 29.08 |
| 5 | Normal | - | - | 22 | M | 170 | 47.70 | 23.76 |
| 6 | Normal | - | - | 22 | M | 170 | 41.34 | 27.28 |
| 7 | Normal | - | - | 52 | M | 188 | 47.22 | 26.33 |
| Mean |  |  |  | 30.1 |  | 171.7 | 43.21 | 28.53 |
| ± SD | - | - | - | ±13.7 | - | ±10.1 | ±3.41 | ±3.13 |

The experimental procedures for this study were consistent with the Ethical Principles for Medical Research Involving Human Participants described in the Declaration of Helsinki and were approved by the Institutional Review Board of Georgia Institute of Technology. NS participants signed the consent form to participate in the study after reading the protocol description. VI participants gave their verbal consent to participate after researchers explained the protocol, read the consent form, and answered questions. A researcher signed and dated the consent form after receiving verbal assent from each VI participant.

### Experimental Tasks

The participants performed bilateral arm matching tasks in the horizontal workspace while seated in the chair of the bimanual Kinarm exoskeleton with two degrees of freedom in each exoskeleton arm for the elbow and shoulder joints (BKIN Technologies Ltd., Kingston ON, Canada). Participants’ pronated arms were supported by exoskeleton arms at the shoulder horizontal level (Fig. 1), and the vertical axis of rotation of the shoulder and elbow joints of each arm aligned with the corresponding Kinarm’s joint axes. No motion at the wrist joints was available. The Kinarm was then calibrated in accordance with established procedures for this exoskeleton (Dukelow et al., 2010).

**Fig. 1.**
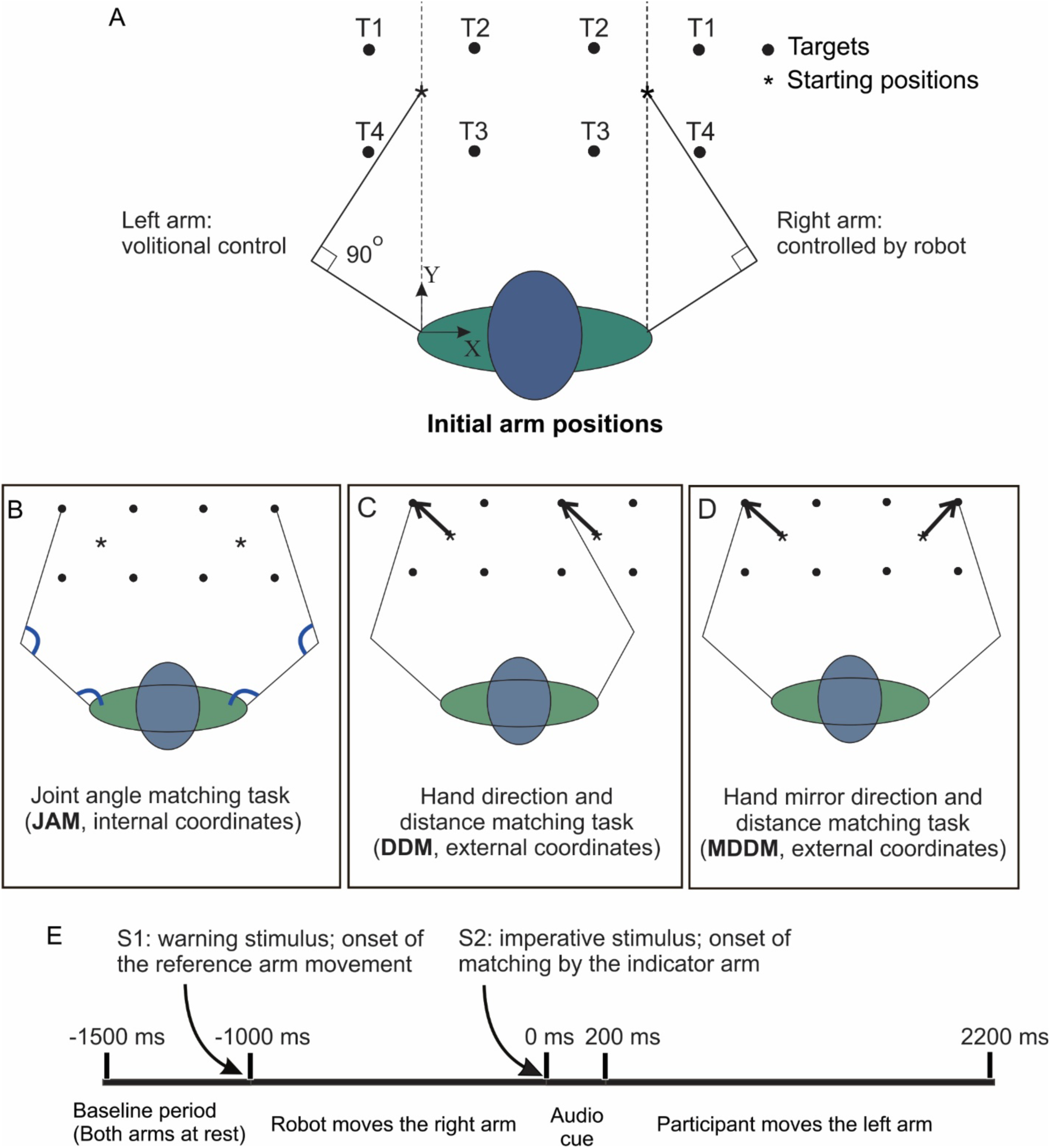
Overview of the experiment and the arm position matching tasks. A: Initial configuration of both the arms at the start of trial. The KINARM robot moves the right arm to one of the four targets in a random order, whereas the left arm is moved under volitional control to match either joint angles or hand location of the right arm. The target locations are marked by black circles and the initial index fingertip positions, by asterisks. B-D: Schematics of the joint angle matching tasks (JAM), hand direction and distance matching task (DDM), and hand mirror direction and distance matching task, respectively. In the JAM task, the robot moves the right hand from the center location to one of the four targets, and the participant is instructed to match the elbow and shoulder joint angles of the right arm with the joint angles of left arm. This arm position matching task is performed in joint angle-based internal coordinates. In the DDM task, the robot moves the right hand to one of the four targets, and the participant is instructed to move their left index fingertip from the initial position in the same direction and to the same distance as the right index fingertip was moved. This hand position matching task is performed in external body-based coordinates. In the MDDM task, the participant is instructed to move their left index fingertip from the initial position in the opposite (mirror) direction and to the same distance as the right index fingertip was moved. This hand position matching task is performed in external body-based coordinates. E: Timeline of events in each experimental trial. Each trial started with a 500-ms period when both arms are stationary with the index fingertips at the initial location as depicted in panel A (-1500 ms to -1000 ms). This period is followed by one second of computer controlled right hand movement to one of the four targets in random order (this period starts with the warning audio stimulus S1 at -1000 ms and ends with the imperative 200-ms audio stimulus S2 at 0 ms). The stimulus S2 is the signal for the participant to start a position matching task with the left arm while the right arm is stationary; the matching task lasts for two seconds and ends at 2200 ms.

At the start of each trial, both participants’ arm endpoints (index fingertips) were placed by the Kinarm at the center of the target workspace such that the index fingertips were directly in front of the corresponding shoulder, and the elbow angles were 90° (Fig. 1A). Each participant performed three bilateral arm matching tasks wherein the robot moved the right (reference) arm to one of 4 targets forming a square in the horizontal workspace and then had to match the reference arm position with their left (indicator) arm. Each participant performed three matching tasks: (1) joint angle matching (JAM, Fig. 1B), in which the participants matched the shoulder and elbow angles of the reference right arm with joint angles of the indicator left arm (matching internal joint coordinates); (2) hand direction and distance matching (DDM, Fig. 1C), in which the participants matched the reference right hand direction and distance from the initial right hand position with the left hand direction and distance from the initial left hand position (matching hand external space coordinates); and (3) hand mirror direction and distance matching (MDDM, Fig. 1D), in which the participants matched the reference right hand mirror direction and distance from the initial right hand position with the left hand mirror direction and distance from the initial left hand position (matching hand external space coordinates). This task was identical to that in our previous study (Oh and Prilutsky, 2022). Note that tasks JAM and MDDM were kinematically identical but differed in the coordinate frames used for matching, i.e., internal in JAM and external in MDDM. In all three matching tasks, the reference right arm of the participant was under computer control while the indicator left arm was under volitional control. The arm matching tasks were performed in following order: JAM, DDM, and MDDM. Each matching task consisted of 72 trials in total, i.e., 18 matchings for each of 4 targets with a 5-min break after every block of 24 repetitions. The order of targets was random.

At the beginning of each trial, the hands of both arms were set in the initial position in front of the corresponding shoulder (Fig. 1A). After a delay of 500 ms, a warning beep S1 indicated that Kinarm started moving the right reference arm to one of the four target positions in a random order; this movement took 1 s. When the Kinarm robot placed the right index fingertip of the arm at a target and stopped, it generated an auditory cue (200-ms beep, S2) indicating that the participant could start moving the indicator left arm to match joint angles or hand horizontal position. The assigned time for performing the matching task was 2.2 s. The participants were instructed to maintain the matched target position of the left arm steady until Kinarm returned the two arms in the initial condition. The sequence of events in each trial is detailed in Fig. 1E. During the experiment, the arms and targets were hidden from the participant’s view by a non-transparent screen and cloth.

In a planar two-joint arm, each combination of shoulder and elbow angles uniquely determines endpoint position. Thus, errors in joint configuration can be equivalently expressed as endpoint spatial errors; e.g. (Oh and Prilutsky, 2022). Therefore, if the left and right joint angles were perfectly matched in the JAM task, the left index fingertip would be at the same relative position as the right index fingertip. The same holds for the MDDM task—a perfect match of the Cartesian coordinates of the right index fingertip by the left fingertip would correspond to a perfect match of angles at the elbow and shoulder joints.

To account for the difference in participants’ arm length, the target distance for each participant (*D_T_*) was calculated based on the distance between the initial index fingertip position and the shoulder: 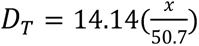, where *x* is the distance in cm between the initial index fingertip position and the shoulder of a given participant; 50.7 is the same distance in cm for a participant from a preliminary study. The lengths of forearm+hand and upper arm of that participant were 40.86 cm and 30.01 cm, respectively. For that subject the distance from the initial index fingertip position to any of the four targets was 14.14 cm, and the targets formed a square with the sides of 20 cm. In this study, the range of target distances was between 11.5 cm and 16.6 cm.

### Kinematic Data Analysis

As mentioned above, any combination of elbow and shoulder angles of a two-joint arm in a horizontal plane uniquely corresponds to the two Cartesian coordinates of the index fingertip. Therefore, to evaluate the accuracy and precision (random errors) of the joint angle matching performance in the JAM task, we analyzed the matching position errors of the indicator left index fingertip in the left workspace with respect to the position of the corresponding reference right index fingertip in the right workspace. The performance of hand position matching in the DDM and MDDM tasks was evaluated by comparing the matching position of the left index fingertip with the corresponding target positions in the left workspace.

The accuracy of position sense (mean error) for each participant, task, and target was computed as the mean distance between the matching position of the left index fingertip and the corresponding required target position in the DDM and MDDM tasks (Fig. 1C-D) or right index location for the JAM task (Fig. 1B). To compute the random error (precision) of the hand position sense, the mean x and y coordinates of the left fingertip obtained across 18 trials for each target, task, and participant were subtracted from the x and y coordinates of the left fingertip in individual trials of the corresponding target, task, and participant. These differences represent random errors of the hand or joint angles position sense. The experimental distribution of these errors was fitted by the 95% confidence ellipse (Johnson and Wichern, 2007), referred to as the precision ellipse in the subsequent text. Ellipse characteristics were quantified by four parameters: the length and orientation of the major axis, area, and shape (ratio of the lengths of the major and minor axes). The ellipse orientation was defined as an angle between its major axis and the horizontal in the range of 0° to 90°. Positive and negative orientation angles of the precision ellipse corresponded to the counterclockwise and clockwise rotations of the major axis from the horizontal, respectively.

### EEG recordings and analyses

Electroencephalography (EEG) was recorded while the participants were performing the hand position or joint angle matching tasks. The participants were wearing a standard 64 channel tin electrode cap (Electro cap, Eaton, OH), and EEG was recorded with Synamps 2 system (Neuroscan, Charlotte, NC). Two electrodes were placed around the left eye to measure the electrooculographic activity, and one electrode was attached at each ear as a linked-ears reference. The sampling rate of the recording was 1000 Hz.

To isolate the CNV potential, raw EEG data were bandpass filtered between 0.5 Hz and 50 Hz. Given that the CNV potential is a slow-wave shift (Tecce, 1972), the lower cutoff frequency was chosen to maintain signal integrity while removing DC drift. Epochs were extracted from -1500 ms to 2200 ms relative to the onset of the imperative stimulus (audio beep S2; Fig. 1E). A baseline period was defined from -1500 ms to -1000 ms (prior to the warning stimulus S1), representing a state where both arms were stationary at the initial position. All subsequent EEG amplitude measurements were calculated relative to the mean voltage of this baseline.

The EEG potential was quantified during two primary phases of interest. (1) Passive/Preparatory Phase (-1000 ms to 0 ms): This window covers the interval between the warning signal (S1) and the imperative signal (S2), after which a participant starts a matching task with the left indicator arm. A slow upward deflection of EEG potential from the baseline between the stimuli S1 and S2 has been associated with the CNV potential (Walter et al., 1964; Tecce, 1972). During this period in our experiments, the robot moved the reference arm, while the left indicator arm was stationary. (2) Active Execution Phase (0 ms to 2200 ms): This window covers the volitional matching movement by the left indicator arm. To capture the temporal evolution of the EEG, the signal was segmented into consecutive 200 ms non-overlapping bins. The mean magnitude within each bin was calculated for each participant, task, and target. Eye blink and movement artifacts in the EEG signal were corrected via recursive least squares (Natraj et al., 2018), and the residual artifacts were corrected with independent component analyses (Natraj et al., 2013; Natraj et al., 2018).

We restricted our analysis of EEG and specifically the CNV potential primarily to the premotor, motor, and centroparietal electrodes over both left (C3A, C3, C3P) and right hemispheres (C4A, C4, C4P), the cortical areas associated with motor planning and spatial integration. The signals within each hemisphere were averaged to provide a robust estimate of the hemispheric-specific EEG activity, including cognitive-motor load.

### Statistics

To test the effects of vision, arm matching task, and target on the accuracy and precision characteristics of hand position sense, a linear mixed model analysis (West et al., 2022) was performed using statistical software IBM SPSS v21 (Chicago, IL, USA). In this analysis, Vision (normal sight, visually-impaired), Task (JAM, DDM, MDDM), and Target (target locations T1 – T4, Fig. 1A) were within-participant independent fixed factors, whereas Participant and Trial were random factors. The dependent variables were accuracy (the mean errors) and precision (random errors, i.e., area of the precision ellipse) in arm matching tasks.

The effects of fixed factors Vision and Task on the mean EEG voltage magnitude within each 200-ms bin across four targets for the right and left hemispheres were tested by a two-way ANOVA (IBM SPSS v21; Chicago, IL, USA). Descriptive statistics values are reported as mean ± 95%-confidence intervals (CI), unless indicated otherwise.

The Bonferroni post-hoc test was used for pairwise comparisons. The significance level for all tests was set at α = 0.05.

## RESULTS

The target positions for each subject were determined by accounting for each participant’s arm lengths (see Method section). However, to display results of accuracy and precision pooled across multiple subjects on the same plot, the coordinates of the fingertip and targets were adjusted so that the distance from the initial fingertip position to each of the four targets was 14.14 cm, i.e. the targets for the right and left arms formed a square with 20-cm sides.

### Accuracy of arm position sense

The mean matching positions of the left index fingertip with respect to the target positions of individual NS and VI participants are shown in Fig. 2. The mean accuracy of hand or joint angle position sense (the distance between the orange and the corresponding black circles) was markedly better in tasks JAM and MDDM than in task DDM in both groups of participants. The accuracy position sense was significantly affected by all three independent factors: Vision (F_1,2608_=94.467, p<0.001), Task (F_2, 2608_=454.817, p<0.001), and Target (F_3, 2608_=27.025, p<0.001). All interactions between these factors were significant (F_2-6,2608_ = 9.141-78.941, p < 0.001).

**Fig. 2.**
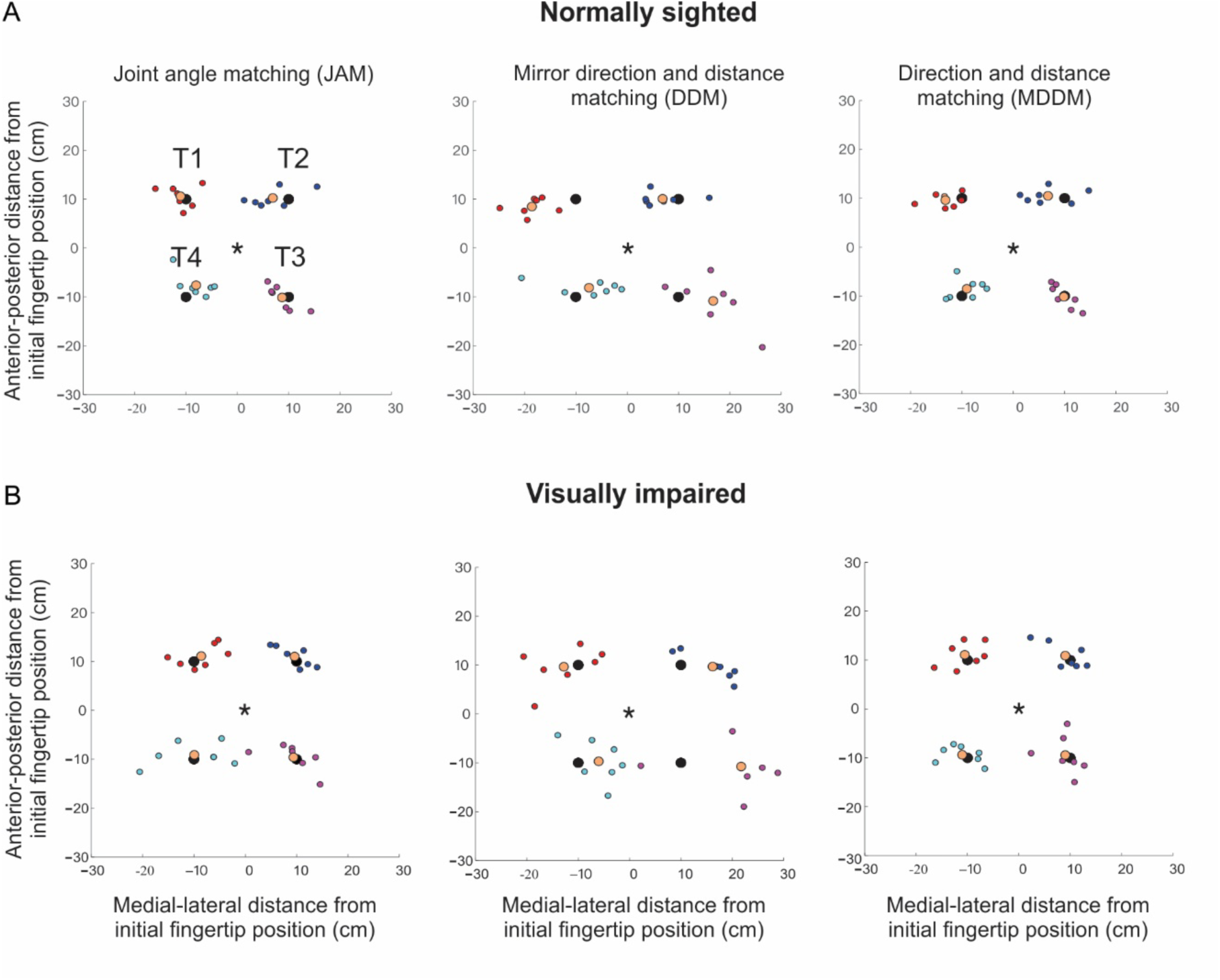
Mean accuracy and precision of arm position matching of individual normally sighted and visually impaired participants in three tasks. Target positions (index fingertip of the right reference arm) are denoted by large black circles. The mean locations of the left index fingertip of the indicator arm a group of participants are denoted by the large orange circles. The mean locations of the left index fingertip of individual participants are denoted by small circles in target-related colors. A: Results of normally sighted participants in three arm position matching tasks. B: Results of visually impaired participants in three arm position matching tasks.

Post-hoc analyses revealed that the accuracy of position sense in all three tasks was better in the NS group compared to the VI participants across all targets (the accuracy is better if the mean deviation from the target, accuracy error, is smaller; Fig. 3A); JAM: F_1, 2608_=19.414, p<0.001; DDM: F_1, 2608_=80.880, p<0.001; MDDM: F_1, 2608_=11.541, p<0.001 (Fig 3A). Another post-hoc analysis for task comparisons revealed that the accuracy of position sense was significantly lower in the DDM task than in the JAM and MDDM tasks in both groups (Fig. 3B); NS: F_2, 2608_=178.585 p<0.001; VI: F_2, 2608_=279.177 p<0.001. No significant difference in accuracy was found between tasks JAM and MDDM in either NS (p=1.000) or VI (p=0.310) participants (Fig 3B).

**Fig. 3.**
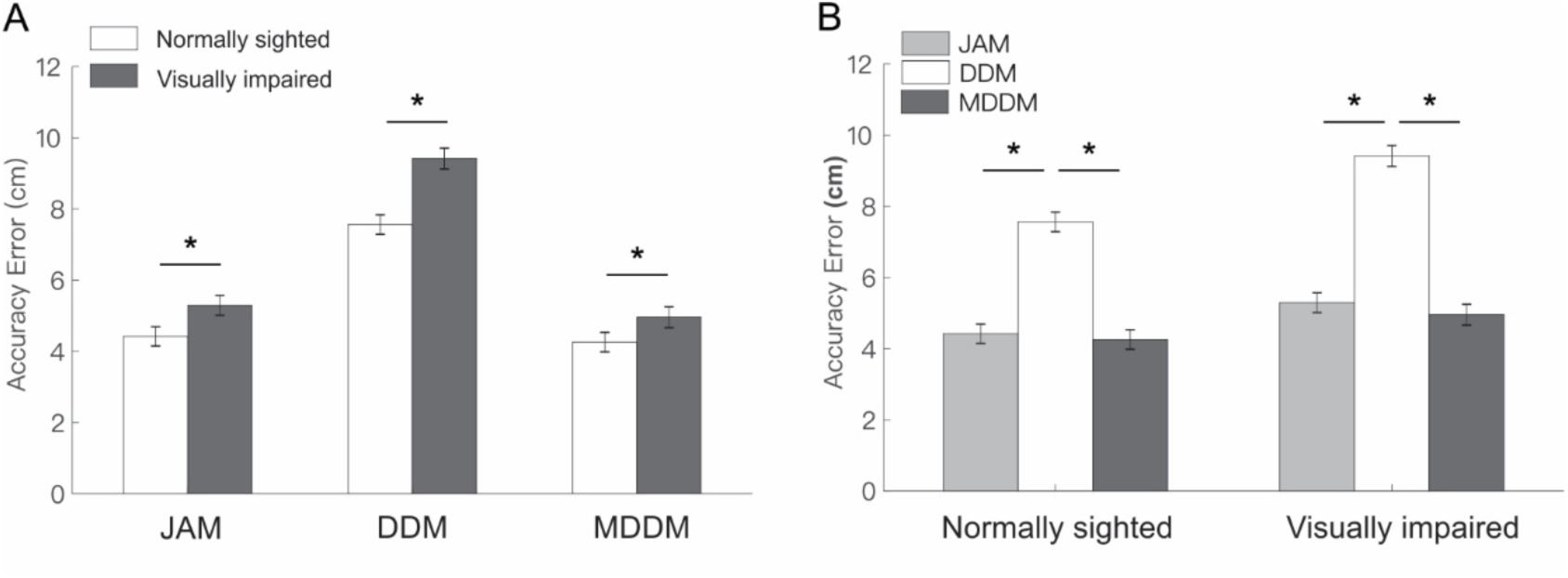
Mean (±CI) distance from the target in the normally sighted and visually impaired groups during three arm position matching tasks. Horizontal lines with an asterisk indicate significant difference (p < 0.05). JAM, DDM and MDDM denote the joint angle matching, hand distance and direction matching, and hand distance and mirror direction matching tasks, respectively. A: The comparison of the mean distance from the targets between normally sighted and visually impaired individuals in three arm position matching tasks. B: The comparison of the mean distance from the target between three arm position matching tasks in normally sighted and visually impaired individuals.

### Precision of arm position sense

The precision (random error) of the arm position sense, as measured by the area of a precision ellipse, was significantly affected by all three independent factors: Vision (F_1,144_=19.902, p<0.001), Task (F_2,144_=5.162, p=0.007), and Target (F_3,144_=2.794, p=0.043). No significant interaction was found between these factors. Precision ellipses of individual participants for each matching task are shown in Fig. 4. Precision ellipses at left targets T1 and T4 typically had positive or horizontal orientations as opposed to ellipses at right targets T2 and T3 that often had negative orientations. Two congenitally blind participants, P1 and P2, as well as VI participant P3 who was blind for 43 years (Table 1), demonstrated the lowest precision of hand position sense in all the tasks (ellipse areas ranged from 22.0 cm^2^ to 244.5 cm^2^; Fig. 4B).

**Fig. 4.**
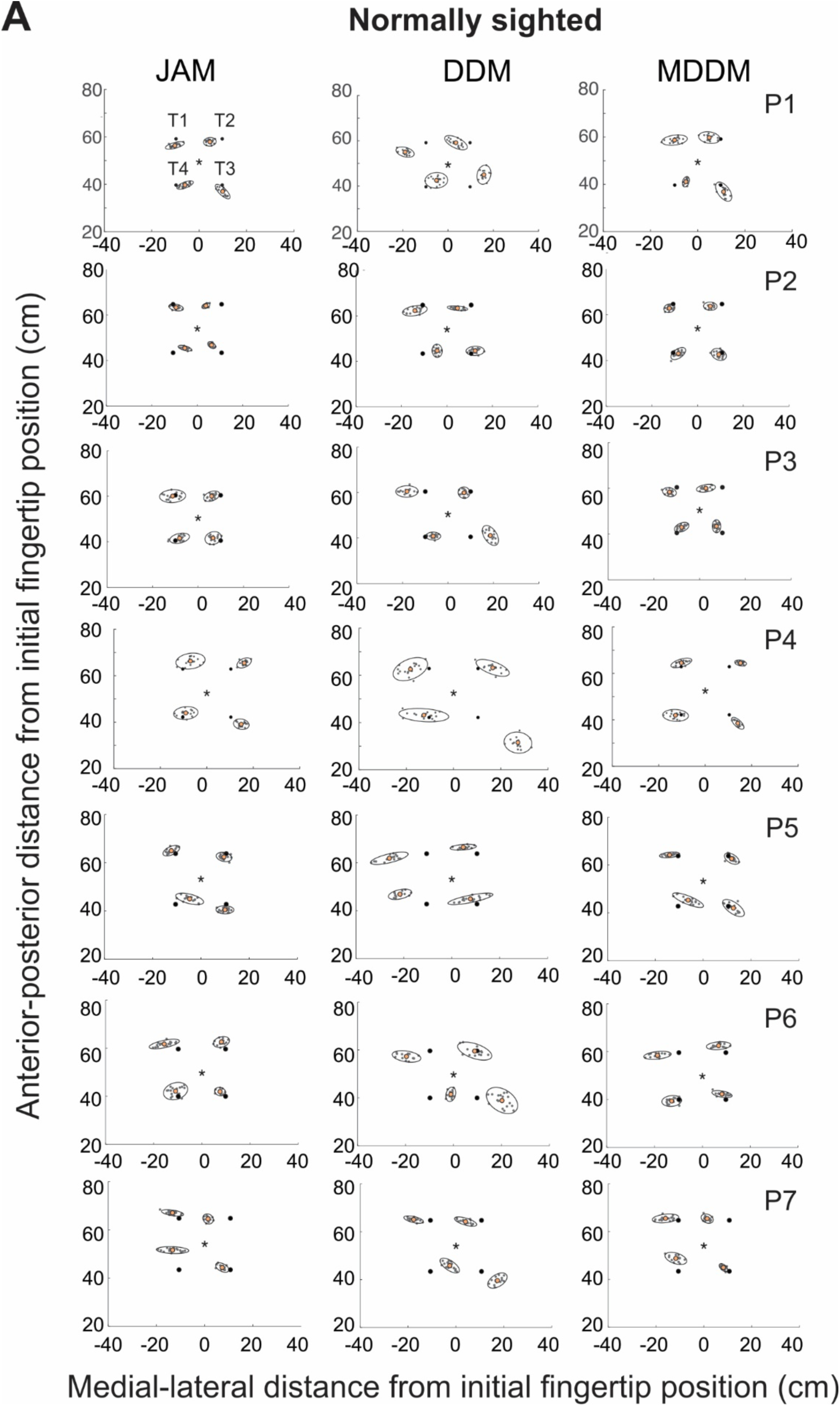
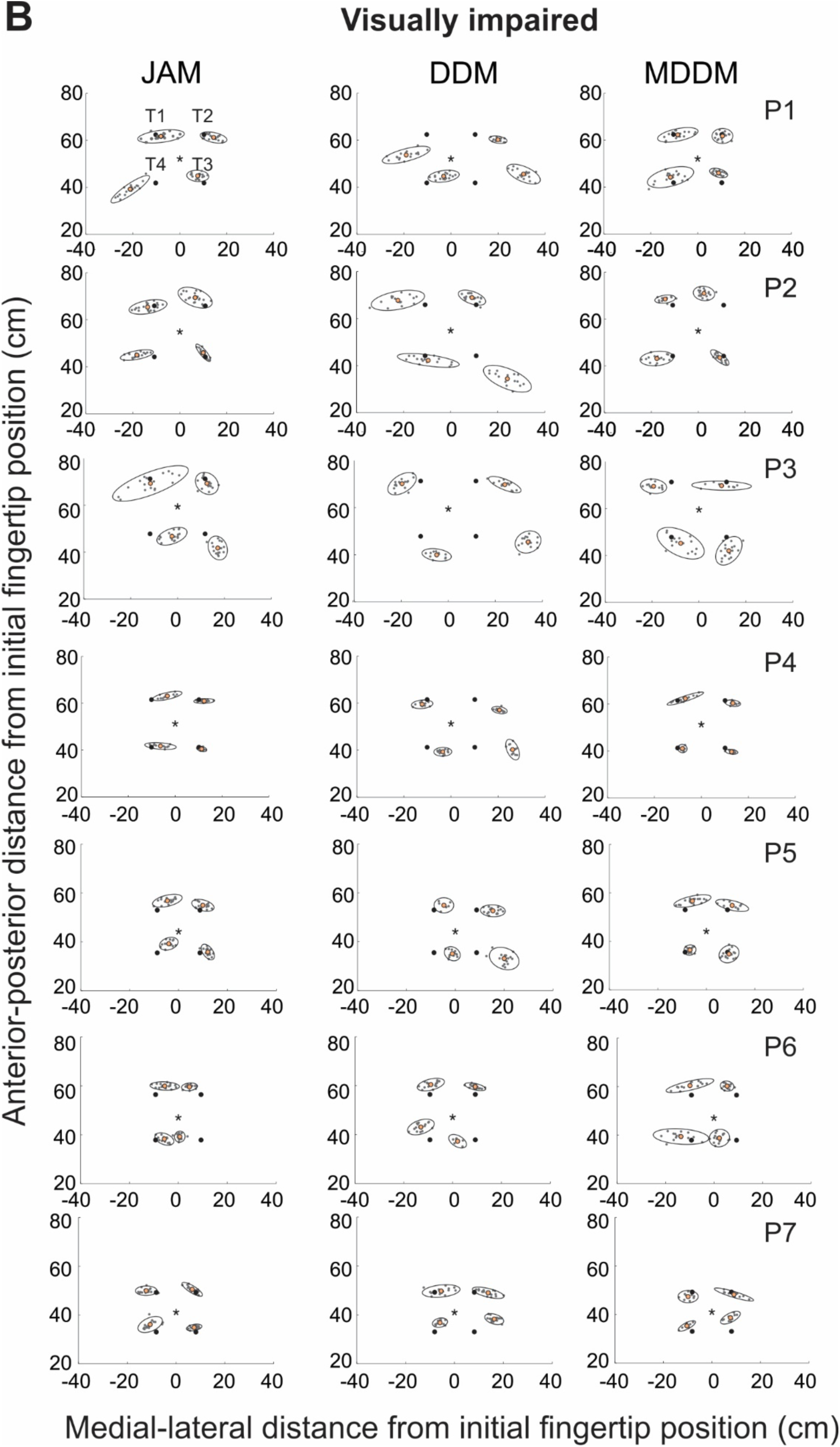
Precision 95%-confidence ellipses of arm position sense for individual participants in three arm position matching tasks. The center of each precision ellipse corresponds to the mean location of the index fingertip of the left indicator hand denoted by the orange circle, whereas gray dots in each ellipse indicate left index fingertip position in individual trials of each participant. Target locations are denoted by the black circles, and fingertip initial position, by the asterisk. JAM, DDM and MDDM denote the joint angle matching, hand distance and direction matching, and hand distance and mirror direction matching tasks, respectively. A: Precision ellipses of seven (P1, P2, …, P7) normally sighted participants. B: Precision ellipses of seven visually impaired participants.

Precision ellipses combined for all participants within the NS and VI groups are shown for each task in Fig. 5. For plotting purposes, the mean positions of the left index finger of each participant were placed to corresponding target locations. Precision ellipses of the VI participants were qualitatively much larger than those of the NS participants in all three tasks. Ellipse orientation for a given target was generally consistent across the tasks for both groups of participants with few exceptions. For example, in VI participants, ellipse orientation at target T3 was positive for the MDDM task but negative in the JAM and DDM tasks (Fig. 5B). Post-hoc analysis revealed higher precision of arm position sense (lower area of the precision ellipse) in the NS group compared to the VI participants in tasks JAM (F_1, 144_ =7.856, p=0.006) and MDDM (F_1, 144_ =11.580, p<0.001). However, there was no difference in precision between the two groups of participants in the DDM task (F_1, 144_ =2.314, p=0.130; Fig. 6A). When the area of the precision ellipses was compared among tasks within each group of participants, the NS group demonstrated lower precision of position sense in the DDM task compared to the JAM (p<0.020) and MDDM (p<0.021) tasks. There was no difference in the precision ellipse area between the JAM and MDDM tasks (p=1.000; Fig. 6B). The precision ellipse areas in the VI group did not differ significantly among the three tasks (F_2, 144_ =1.082, p=0.342; Fig. 6B).

**Fig. 5:**
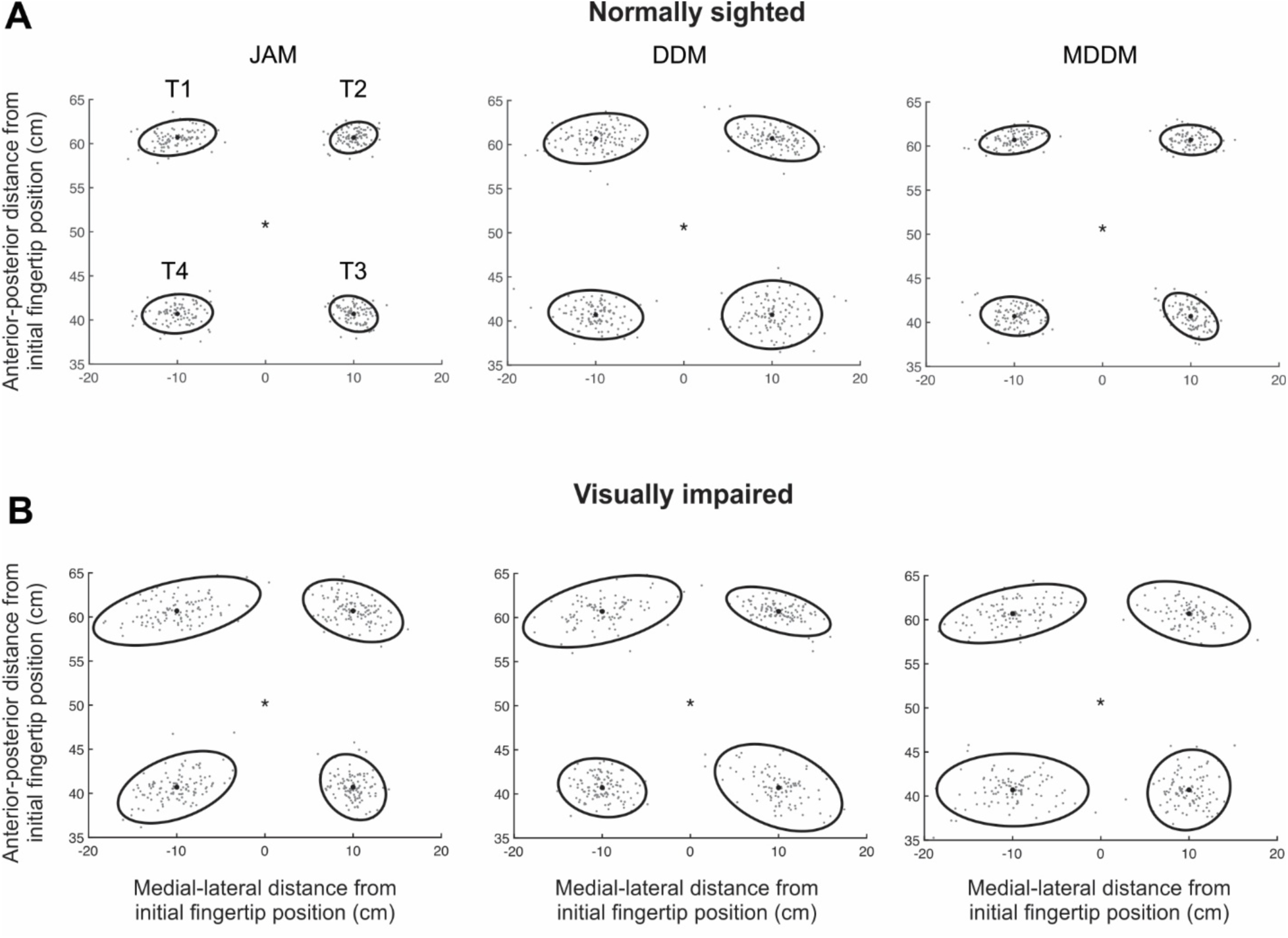
Precision 95%-confidence ellipses of arm position sense averaged across participants within the normally sighted and visually impaired groups for each arm position matching task. For display purposes, the mean position of the index fingertip of the left indicator hand of each participant is placed at the corresponding target location. JAM, DDM and MDDM denote the joint angle matching, hand distance and direction matching, and hand distance and mirror direction matching tasks, respectively. A: The mean precision ellipses of normally sighted participants. B: The mean precision ellipses of visually impaired participants.

**Fig. 6.**
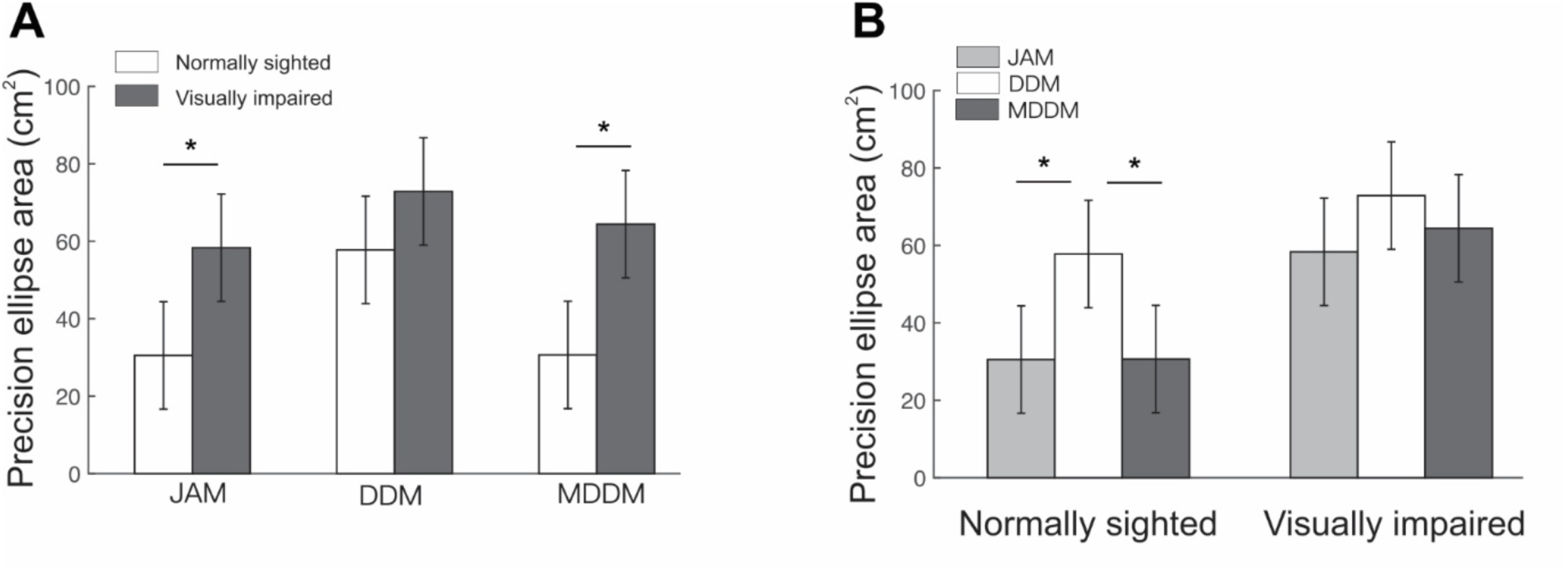
Mean (±CI) precision ellipse area in the normally sighted and visually impaired groups during three arm position matching tasks. Horizontal lines with an asterisk indicate significant difference (p < 0.05). JAM, DDM and MDDM denote the joint angle matching, hand distance and direction matching, and hand distance and mirror direction matching tasks, respectively. A: The comparison of the mean precision ellipse area between normally sighted and visually impaired individuals in three arm position matching tasks. B: The comparison of the mean precision ellipse area between three arm position matching tasks in normally sighted and visually impaired individuals.

### EEG activity and CNV potential

The mean cortical head maps of EEG activity at four specific snapshots through the trial (-400 ms, -50 ms, 450 ms, and 700 ms) for the two participant groups and across the three tasks are shown in Fig. 7A-D. The first two time instants, -400 ms and -50 ms, were within the warning interval (S1-S2: -1000 ms – 0 ms), during which participants prepared to initiate their motor response, i.e., an active arm position matching by the left indicator arm. The initial time instant, - 400 ms, occurred after a typical delay before developing the CNV, while the second, -50 ms, was just before the expected imperative stimulus S2 and during a decline of the CNV potential (Tecce, 1972). At -400 ms, there were elevated negative potentials in the central region covering the sensorimotor and parietal cortices in both groups of subjects. The VI group also demonstrated greater negativity in the frontal areas and positive potentials in the occipital cortex. There was a trend of increasing the negative potentials around the apex of the skull in the NS groups from JAM to MDDM and to DDM tasks; the opposite trend occurred in the VI group, i.e., the negative potentials decreased from JAM to MDDM and to DDM tasks (Fig. 7A). At -50 ms, the negative potentials occurred in the same central areas, although their area and magnitude increased in the VI group from the lowest values in the JAM task to the highest values in the DDM task (Fig. 7B).

**Fig. 7:**
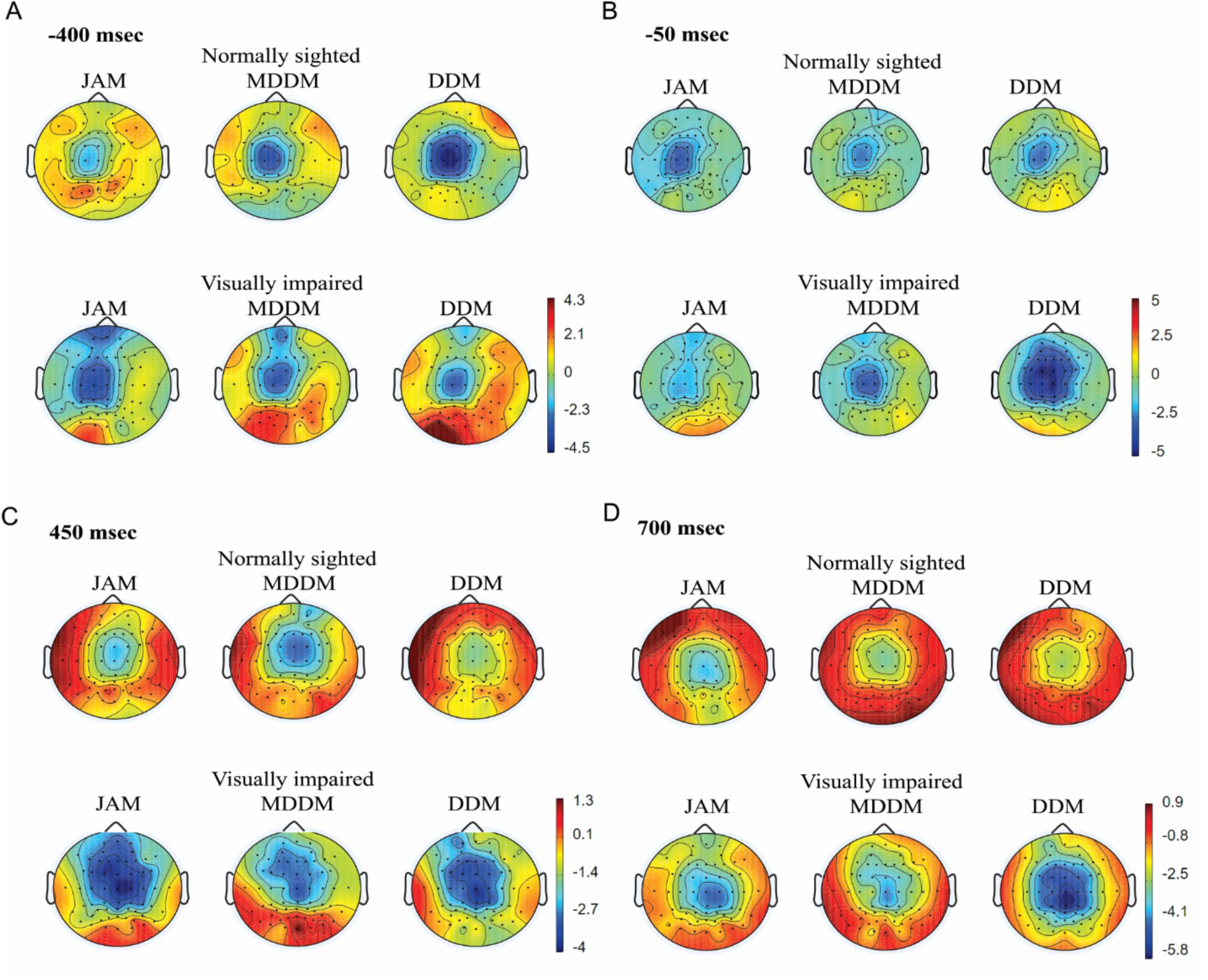
Mean cortical heat maps of EEG activity at specific time snapshots. Cortical activity projected on a 2D surface representing the EEG grid through the trial. Individual dots represent electrodes. Color represents EEG field potential values. Data is shown for both participant groups (rows) and for each of the three tasks (columns). JAM, DDM and MDDM denote the joint angle matching, hand distance and direction matching, and hand distance and mirror direction matching tasks, respectively. A: EEG activity at -400 ms when the right arm was being moved by the KINARM robot and the left arm was stationary. B: EEG activity at -50 ms when the right arm was being moved by the KINARM robot and the left arm was stationary. C: EEG activity at 450 ms when the right arm was stationary while the left arm performed a matching task. D: EEG activity at 700 ms when the right arm was stationary while the left arm performed a matching task.

The two last time snapshots depicted in Fig. 7 (450 ms and 700 ms) occurred after the imperative stimulus S2 and the onset of the motor response, i.e., the movement of the left indicator arm towards the target matching position. Similar to the CNV potential, the central regions had negative or lower positive potentials compared to the other head regions in both groups of participants and both time snapshots (Fig. 7C, D). The VI group had generally greater negative EEG magnitudes, and the negative shift occurred also in the frontal areas, especially at 450 ms. The VI group demonstrated a much greater negative potential in task DDM at 700 ms compared to the other tasks of this group. Thus, the CNV potential in the sensorimotor and parietal cortices could be seen in both groups of participants during the waiting period (S1-S2).

We then compared the cortical potentials averaged across the premotor, motor and centroparietal electrodes for the left (C3A, C3, C3P) and right (C4A, C4, C4P) hemispheres separately. These mean EEG activity patterns for the NS and VI groups for each arm position matching task are shown in Fig. 8A-C for the left hemisphere and in Fig. 8D-F for the right hemisphere. Qualitatively, the CNV (a negative potential shift during the waiting period) is clearly seen in both groups of participants and the three tasks starting between -500 ms and -400 ms and lasting until the imperative stimulus S2 at 0 ms. During the imperative audio stimulus S2 (0 to 200 ms), there was a sharp increase in the negative potential followed by its quick decrease to positive values in the two groups of participants and the three tasks. Subsequently, there was a negative shift of the mean EEG potential again that typically lasted beyond 1000 ms. Although the VI group appeared to produce larger negative shifts with respect to the baseline before and after stimulus S2 than the NS group in both hemispheres and the tasks JAM and DDM, a statistical difference was found only in the period -400 ms - -200 ms in the left hemisphere and the task JAM (F_1,12_ = 1.251, p=0.023; Fig. 8A) and the period 600 ms – 800 ms of the same hemisphere and the task DDM (F_1,12_ = 0.190, p=0.020; Fig. 8B).

**Fig. 8:**
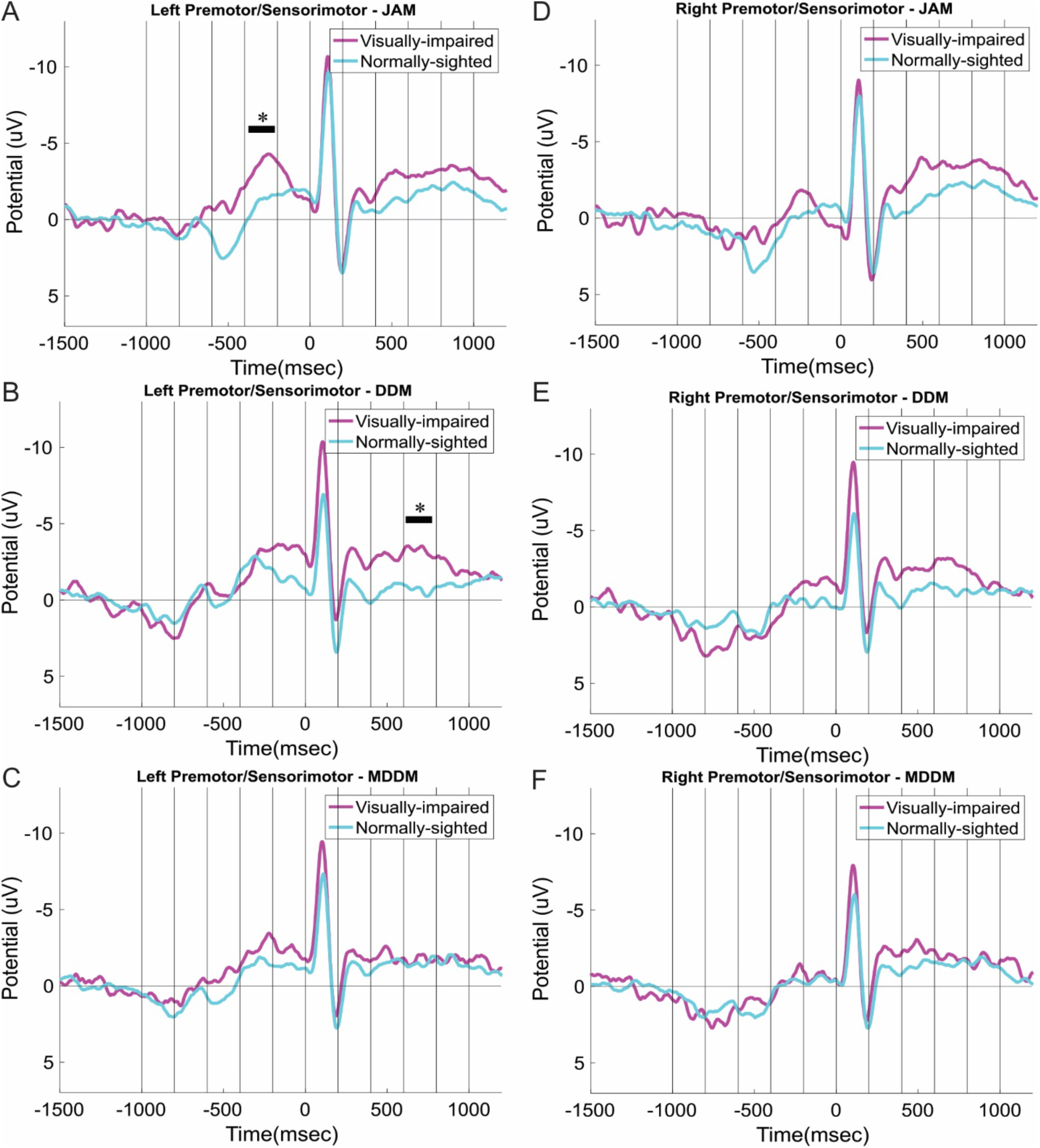
Comparisons of the mean premotor/sensorimotor EEG potential in the left hemisphere (left column) and right (right column) hemisphere between the normally sighted and visually impaired groups during three arm position matching tasks (rows). The EEG potentials in the left column were obtained by averaging the combined activity recorded by electrodes C3A, C3, and C3P in all trials and participants within a given position matching task and the participant group. The EEG potentials in the right column were obtained in the same way except recordings of electrodes C4A, C4, and C4P were used. Thick horizontal lines with asterisk indicate significant difference in EEG potential during the corresponding 200-ms time bin between normally sighted and visually impaired participants. The horizontal time axis corresponds to the time axis in Fig. 1E. A-C: EEG potentials for the left premotor-sensorimotor cortex in JAM, DDM, and MDDM position matching tasks, respectively. D-F: EEG potentials for the right premotor-sensorimotor cortex in JAM, DDM, and MDDM position matching tasks, respectively.

Comparing the mean cortical potentials between the various arm matching tasks within each participant group and hemisphere (Fig. 9), we found that the NS participants had a greater CNV potential in the left hemisphere in the DDM task compared to the JAM task during the waiting period between -600 ms and -400 ms (p=0.023) and between -400 ms and -200 ms (p=0.043); similar significant difference was obtained after the imperative stimulus at -800 ms -1000 ms (p=0.007; Fig. 9A). The NS participants did not demonstrate a typical CNV potential during the waiting period in the right hemisphere; instead, the mean cortical potentials were positive or close to the baseline for all three tasks, and the positive potential in JAM task was statistically greater than that in the DDM task at the time window -600 ms - -400 ms (Fig. 9B). In the end of trial (800 ms – 1000 ms) of the NS group, the negative cortical potential in the right hemisphere was greater in the JAM task than in the DDM task (p=0.005; Fig. 9B).

**Fig. 9:**
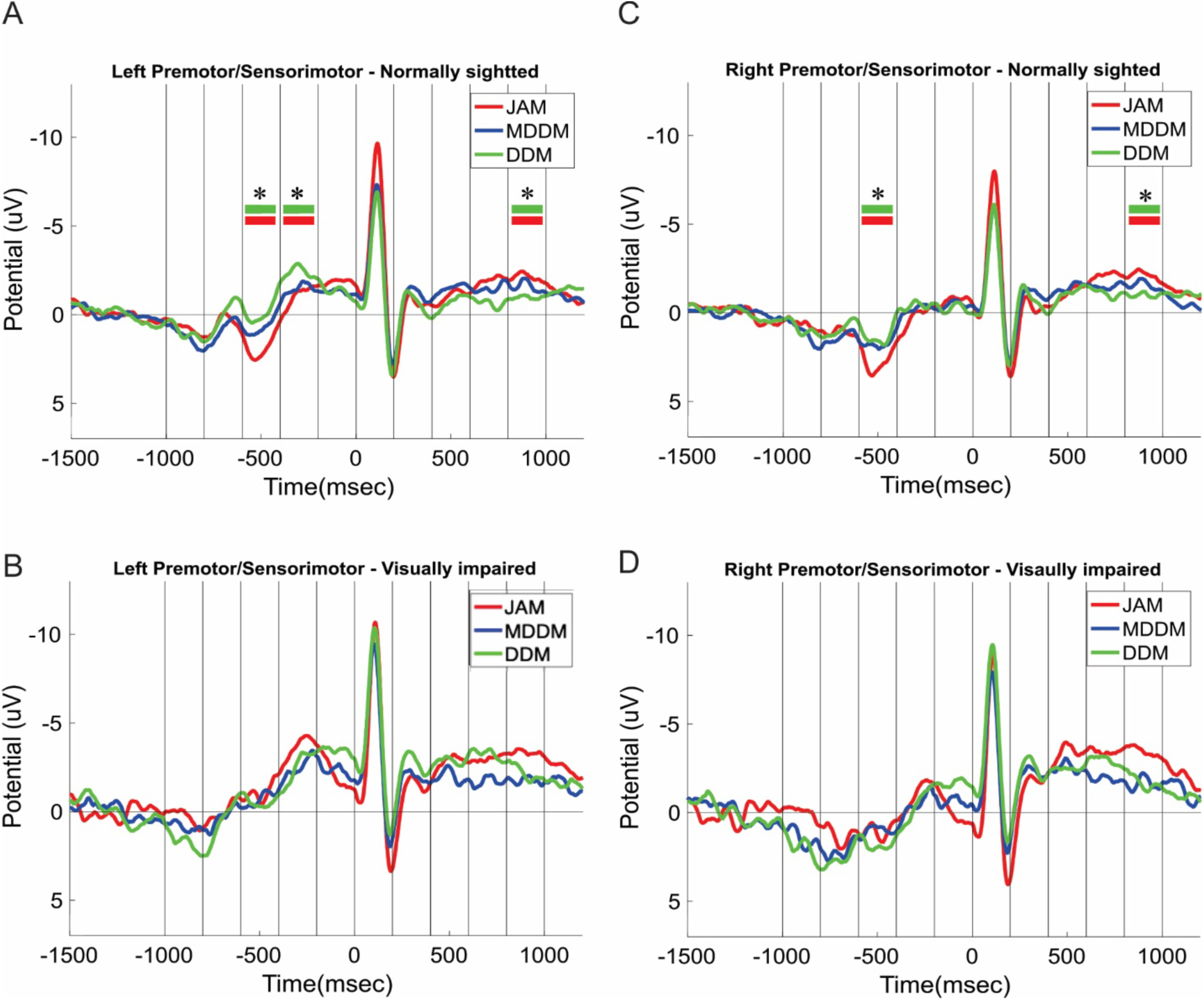
Comparisons of the mean premotor/sensorimotor EEG potential in the left hemisphere (left column) and right (right column) hemisphere among three arm position matching tasks for the normally sighted and visually impaired groups (rows). The EEG potentials in the left column and right column were obtained in the same way as in Fig. 8. Thick green and red horizontal lines with asterisk indicate significant difference in EEG potential during the corresponding 200-ms time bin between the JAM and DDM tasks. The horizontal time axis corresponds to the time axis in Fig. 1E. A, B: EEG potentials for the left premotor-sensorimotor cortex in position matching tasks JAM, DDM, and MDDM for normally sighted and visually impaired participants, respectively. C, D: EEG potentials for the right premotor-sensorimotor cortex in position matching tasks JAM, DDM, and MDDM for normally sighted and visually impaired participants, respectively.

The VI participants demonstrated a clear CNV potential in the left hemisphere during the waiting period and a negative potential shift after the imperative stimulus S2. However, no differences were found in the cortical potentials between the matching tasks (Fig. 9C). For the right hemisphere in the VI group, the results were generally similar except there was no discernable CNV potential during the waiting period (Fig. 9D).

## Discussion

The present study examined the effects of visual experience on arm position sense and its associated cortical activity across tasks that required either internal (joint-based) or external (endpoint-based) representations of limb position. Three main findings emerged. First, VI participants exhibited reduced accuracy and, in most conditions, reduced precision of arm position sense compared with NS participants. Second, performance in tasks requiring external coordinate representations (DDM) was consistently worse than in tasks compatible with joint-based representations (JAM), irrespective of visual status. Third, increased cognitive-motor load, denoted by the contingent negative variation (CNV), was associated with task demands and was generally greater in the VI group and in tasks requiring transformations to external coordinates. Together, these results indicate that visual experience contributes to the calibration of arm position sense and that transformations from joint-based to external representations impose additional computational demands.

### Visual experience and position sense

VI participants demonstrated lower accuracy across all matching tasks and reduced precision in JAM and MDDM. These findings suggest that visual experience contributes not only to external spatial representations, but also to the fidelity of internal representations of limb configuration. Although proprioceptive and cutaneous afferents provide the primary signals for joint-based position sense (Proske and Gandevia, 2018; Banks et al., 2021), visual input likely plays a critical role during development in calibrating these signals and establishing accurate body representations (Thinus-Blanc and Gaunet, 1997; Crollen and Collignon, 2012).

The absence of a precision deficit in the DDM task between groups suggests that task constraints can limit the contribution of visual experience. In DDM, both groups exhibited similarly low precision, indicating that the transformation to external coordinates may introduce variability that dominates group differences. Thus, the influence of visual experience appears to be most evident when task demands allow accurate use of internal representations.

These findings are consistent with the view that vision serves as a calibration signal for multisensory integration underlying body schema (Mon-Williams et al., 1997; Nava et al., 2014). Reduced visual experience may lead to less accurate mappings between joint configurations and external space, as well as increased uncertainty in these mappings (Fiehler et al., 2009; Cappagli et al., 2017).

### Internal versus external representations

A central result of this study is the dissociation between tasks requiring explicit transformation to external coordinates (DDM) and those that do not (JAM), as well as the unexpected similarity between JAM and MDDM. Although MDDM was defined in external coordinates, its kinematic equivalence to JAM allowed it to be solved using joint-based representations. The comparable accuracy, precision, and CNV magnitude between these two tasks strongly suggest that participants preferentially relied on internal representations when possible.

In contrast, the DDM task required a transformation from joint angles to endpoint displacement that differed between arms, preventing the use of a simple mirror mapping. This requirement resulted in decreased accuracy and increased variability in both groups. These findings support the hypothesis that computing limb position in external coordinates introduces additional sources of error, likely due to nonlinear transformations and uncertainty in limb geometry (Soechting and Terzuolo, 1986; Longo and Haggard, 2010; Oh and Prilutsky, 2022).

Importantly, these results indicate that differences in performance across tasks cannot be attributed solely to the coordinate frame specified in the instructions, but rather to whether the task permits the use of more direct internal representations. This distinction highlights the importance of considering both computational demands and available sensorimotor strategies when interpreting position-matching behavior.

### Cognitive-motor load and CNV

The CNV results provide insight into the neural correlates of these task-dependent differences. Across conditions, tasks associated with lower behavioral performance (DDM) tended to exhibit larger CNV amplitudes, consistent with increased cognitive-motor demands (Walter et al., 1964; Tecce, 1972). This effect was particularly evident in the left hemisphere, in agreement with its proposed role in sensorimotor integration and spatial transformations in right-handed individuals (Janssen et al., 2011).

VI participants showed a general tendency toward greater negative potentials, suggesting increased neural effort during task preparation and execution. However, statistically significant group differences were limited, indicating variability across participants and conditions. These findings suggest that CNV reflects task difficulty and processing demands rather than group differences per se.

The association between increased CNV magnitude and reduced accuracy supports the interpretation that transforming proprioceptive information into external coordinates requires additional neural resources. At the same time, the absence of strong and consistent statistical differences underscores the need for caution in directly linking CNV amplitude to behavioral performance. Rather, CNV appears to indicate the preparatory and computational demands associated with task execution (Cui et al., 2000).

### Preferred coordinate representation

An important implication of these results is that the motor system may preferentially utilize joint-based representations when they are sufficient for task performance. The similarity between JAM and MDDM indicates that even when instructions emphasize external spatial matching, participants may rely on internal representations if the task structure allows it. This preference likely reflects the reduced computational cost of avoiding coordinate transformations.

Such a strategy would be advantageous given that transformations from joint space to external space are nonlinear and depend on accurate estimates of segment lengths and body configuration (Longo and Haggard, 2010; Naito et al., 2016). Errors in these estimates, particularly in the absence of visual calibration, would propagate into external position estimates. Thus, reliance on joint-based representations may provide a more robust and efficient solution when available.

### Limitations of the study

Several limitations should be acknowledged. First, the sample size was relatively small, which may have reduced statistical power, particularly in the EEG analyses. Second, the VI group included participants with heterogeneous onset and duration of visual impairment, which could differentially affect the development of spatial representations. Congenital and early blindness may allow compensatory plasticity through enhanced use of nonvisual modalities (Lessard et al., 1998; Pascual-Leone et al., 2005), which could partially mitigate deficits in external spatial representation. Third, although MDDM was defined as an external-coordinate task, the possibility that participants solved it using joint-based strategies cannot be excluded.

Future studies could address these issues by increasing sample size, stratifying participants based on onset of visual impairment, and incorporating task designs that more stringently dissociate internal and external representations.

### Conclusions

In summary, visual experience enhances the accuracy and precision of arm position sense and reduces the neural cost of position matching. Tasks requiring transformations to external coordinates degrade performance and increase cognitive-motor load, particularly when joint-based strategies cannot be used. The results further suggest that the sensorimotor system preferentially relies on internal representations when task constraints permit, reflecting an efficient strategy that minimizes computational demands.

## Acknowledgements

This work was supported by the National Science Foundation EFRI-M3C program (Award No. 1137172)

